# The Presence of the Temporal Horn Exacerbates the Vulnerability of Hippocampus during Head Impacts

**DOI:** 10.1101/2021.12.07.471634

**Authors:** Zhou Zhou, Xiaogai Li, August G Domel, Emily L Dennis, Marios Georgiadis, Yuzhe Liu, Samuel J. Raymond, Gerald Grant, Svein Kleiven, David Camarillo, Michael Zeineh

**Affiliations:** Department of Bioengineering, Stanford University, Stanford, CA, 94305, USA; Neuronic Engineering, KTH Royal Institute of Technology, Stockholm, 14152, Sweden; TBI and Concussion Center, Department of Neurology, University of Utah, Salt Lake City, 84108, USA; Department of Radiology, Stanford University, Stanford, CA, 94305, USA; Department of Neurosurgery, Stanford University, Stanford, CA, 94305, USA; Department of Neurology, Stanford University, Stanford, CA, 94305, USA; Department of Mechanical Engineering, Stanford University, Stanford, CA, 94305, USA

**Keywords:** Hippocampal injury, temporal horn, brain-ventricle interface, fluid-structure interaction, finite element analysis, traumatic brain injury

## Abstract

Hippocampal injury is common in traumatic brain injury (TBI) patients, but the underlying pathogenesis remains elusive. In this study, we hypothesize that the presence of the adjacent fluid-containing temporal horn exacerbates the biomechanical vulnerability of the hippocampus. Two finite element models of the human head were used to investigate this hypothesis, one with and one without the temporal horn, and both including a detailed hippocampal subfield delineation. A fluid-structure interaction coupling approach was used to simulate the brain-ventricle interface, in which the intraventricular cerebrospinal fluid was represented by an arbitrary Lagrangian-Eulerian multi-material formation to account for its fluid behavior. By comparing the response of these two models under identical loadings, the model that included the temporal horn predicted increased magnitudes of strain and strain rate in the hippocampus with respect to its counterpart without the temporal horn. This specifically affected cornu ammonis (CA) 1 (CA1), CA2/3, hippocampal tail, subiculum, and the adjacent amygdala and ventral diencephalon. These computational results suggest that the presence of the temporal horn exacerbate the vulnerability of the hippocampus, highlighting the mechanobiological dependency of the hippocampus on the temporal horn.

## Introduction

Traumatic brain injury (TBI) is a critical public health and socio-economic problem. In the United States, approximately 5.3 million people are living with a TBI-related disability (Langlois and Sattin, 2005). At a global level, an estimated 69 million people suffer a TBI each year (Dewan et al., 2018), with yearly costs reaching 400 billion dollars (Maas et al., 2017). Despite worldwide efforts to reduce the incidence and mitigate the consequence of TBI, improvement of overall outcome has not been achieved (Roozenbeek et al., 2013), especially for mild TBI (mTBI), also known as concussion. Epidemiological data showed that concussion rates in high school sports (Rosenthal et al., 2014) and the military (Cameron et al., 2012) have been rising. The need to improve concussion outcome is particularly urgent, given that concussion is notoriously underreported, difficult to screen, and associated with immediate and persistent deficit to memory and attention with possible chronic neurodegenerative consequences (McKee et al., 2015;Meier et al., 2015).

As a crucial structure for long-term, episodic memory formation and retrieval (Bird and Burgess, 2008), the hippocampus is often reported to be injured secondary to physical trauma in humans across different impact severities. In fatal TBI, post-mortem histopathological examinations identify the hippocampus as one of the most commonly injured regions (73%-87%) (Kotapka et al., 1992;Kotapka et al., 1993;Kotapka et al., 1994;Maxwell et al., 2003). In mTBI, *in vivo* human imaging analyses demonstrate that repetitive concussive impacts or even sub-concussive impacts (i.e., high-velocity impacts that do not cause concussion) are associated with abnormal hippocampal atrophy longitudinally (Parivash et al., 2019) and cross-sectionally (Singh et al., 2014). The prevalence of hippocampal injury has also been widely noted in animal experiments (e.g., non-human primates, pigs, rats, sheep, and rabbits) under diverse modes of mechanical perturbations, including non-impact acceleration (Gennarelli et al., 1982;Kotapka et al., 1991), impact acceleration (Anderson et al., 2003), weight-drops (Kalish and Whalen, 2016), cortical contusion (Baldwin et al., 1997), and fluid percussion injury (Hicks et al., 1996). The resultant injury within the hippocampus of experimentally traumatized animals exhibits a broad spectrum of pathological manifestations, varying from impaired electrophysiological activity associated with hippocampal circuitry dysfunction (Wolf et al., 2017) to profound neuronal apoptosis and marked gliosis (Smith et al., 1997).

The pathogenetic mechanism of trauma-induced hippocampal injury has long been attributed to the selective vulnerability of hippocampal neurons to hypoxemia and ischemia (Pulsinelli, 1985;Ng et al., 1989), typical complications of severe TBI insults (Graham et al., 1978;Graham et al., 1989). For example, a histopathological study revealed that 27 out of 29 individuals with at least one episode of clinically recorded hypoxia had hippocampal damage (Kotapka et al., 1992). However, 14 out of 18 patients without documented hypoxemia also had hippocampal lesions (Kotapka et al., 1992), suggesting that hippocampal injury may be independent of hypoxia. Another candidate mechanism is pathological neuronal excitation involving glutamate and/or other excitatory amino acid neurotransmitters, supported by animal experiments where traumatic insults triggered glutamate concentrations in the extracellular fluid of the hippocampus (Faden et al., 1989;Runnerstam et al., 2001). Given that the hippocampus is dense in receptors for glutamate (Kotapka et al., 1991;Leranth et al., 1996), redundant extracellular glutamate could induce neuronal excitotoxicity, and indeed, pre-treatment of experimentally traumatized animals with glutamate antagonists attenuates hippocampal lesions (Faden et al., 1989). However, such antagonists in humans have not proven beneficial, thus, a neuroexcitotoxic mechanism in human TBI cannot be considered a sole explanation (Parsons et al., 1999). Taken together, trauma-induced hippocampal lesions in humans cannot be fully explained by the current mechanisms.

An alternative line of investigation is biomechanical. Given that previous modeling work has shown that the presence of fluid can affect the transmission of mechanical forces within the brain (Zhou et al., 2020a), one structure that may be associated with the hippocampal vulnerability is the temporal horn of the lateral ventricle. The temporal horn is a cavity that forms the roof of the hippocampus and is filled with cerebrospinal fluid (CSF) and occasionally choroid plexus (Insausti and Amaral, 2003). Previous studies found that the volumes of the hippocampus and temporal horn were inversely correlated in TBI patients (Gale et al., 1994;Bigler et al., 1997;Bigler et al., 2002). However, the biomechanical effect of the temporal horn on the hippocampus remains unknown.

Interrogation of this biomechanical relationship requires modeling to estimate the myriad variables and forces at play. As computational surrogates of the human head, finite element (FE) models have been instrumental in exploring the association of regional vulnerabilities with potential predisposing factors during trauma from the biomechanical perspective (Kleiven, 2007;McAllister et al., 2012;Mao et al., 2013;Ji et al., 2015;Atsumi et al., 2018;Trotta et al., 2020;Zhou et al., 2021a). Extending the current models to investigate the relationship between the temporal horn and hippocampus requires that the FE model possesses an anatomically and mechanically accurate representation of both structures, and a precise description of the interface between the fluid-filled temporal horn and neighboring hippocampus. However, in existing finite element models, the temporal horn was either wholly substituted as brain parenchyma (McAllister et al., 2012;Zhou et al., 2016) or simulated as a solid structure using the Lagrangian approach (Kleiven, 2007;Mao et al., 2013;Ji et al., 2015;Atsumi et al., 2018;Trotta et al., 2020;Zhou et al., 2021a). This Lagrangian approach is a dominant numerical scheme for solid mechanics and is insufficient to computationally reflect the fact that the temporal horn is filled with CSF with the potential flow within the ventricular cavity during the impacts (Souli et al., 2000;Zhou, 2019;Zhou et al., 2020b). Approaches to date may have missed key and relevant properties of the temporal horn that have precluded the determination of its biomechanical relevance.

The aim of the current study was to discern whether the presence of the temporal horn exacerbates the biomechanical vulnerability of the hippocampus. To test this hypothesis, two models with and without a detailed anatomic description of the temporal horn profiles are established. By comparing the strain-related responses to identical loadings between the two models, the biomechanical mechanism for the temporal horn’s role in the vulnerability of the hippocampus was uncovered.

## Materials and Methods

In this study, we employed computational modeling to discern the biomechanical dependency of the hippocampus on the temporal horn. To achieve that, we utilized a novel, multi-million element 3D head model (Zhou et al., 2020a) that did not initially incorporate the temporal horn (no-temporal-horn (NTH)-model), and further extended this model by adding the temporal horn to the lateral ventricle (temporal-horn (TH)-Model). An arbitrary Lagrangian-Eulerian (ALE) multi-material formation was used to emulate the fluid behavior of the intraventricular CSF, with its responses being concatenated with the brain tissue via a fluid-structure interaction (FSI) coupling algorithm. This allows computation of strain (fractional change in unit length), strain rate (strain change over time), and stress (force per unit area) in the hippocampus. By comparing the deformation-related responses estimated by these two models secondary to six concussive/sub-concussive impacts, the mechanical role that the temporal horn exerted on the hippocampus was revealed.

### Finite element modeling of human brain

The FE head model without the temporal horn (i.e., the NTH-Model) used in this study was previously developed at KTH Royal Institute of Technology in Stockholm, Sweden (Zhou et al., 2020a). The model includes the scalp, skull, brain, subarachnoid CSF (i.e., CSF within the subarachnoid space), meninges (i.e., dura mater and pia mater), falx, tentorium, and cerebral ventricles (i.e., lateral ventricles without the temporal horn, and third ventricle) (Fig. 1). The whole head model consists of 4.2 million hexahedral elements and 0.5 million quadrilateral elements, in which the brain has a total of 2.6 million nodes, and 2.3 million hexahedral elements. The average brain element size is 0.59 ± 0.26 mm, meeting the requirement that a human brain model with converged responses should have an average element size less than 1.8 mm (Zhao and Ji, 2019). Information regarding the geometry profiles and material modeling of various intracranial components in the NTH-Model was elaborated in a previous study (Zhou et al., 2020a) as well as in Appendix A.

**Fig. 1.**
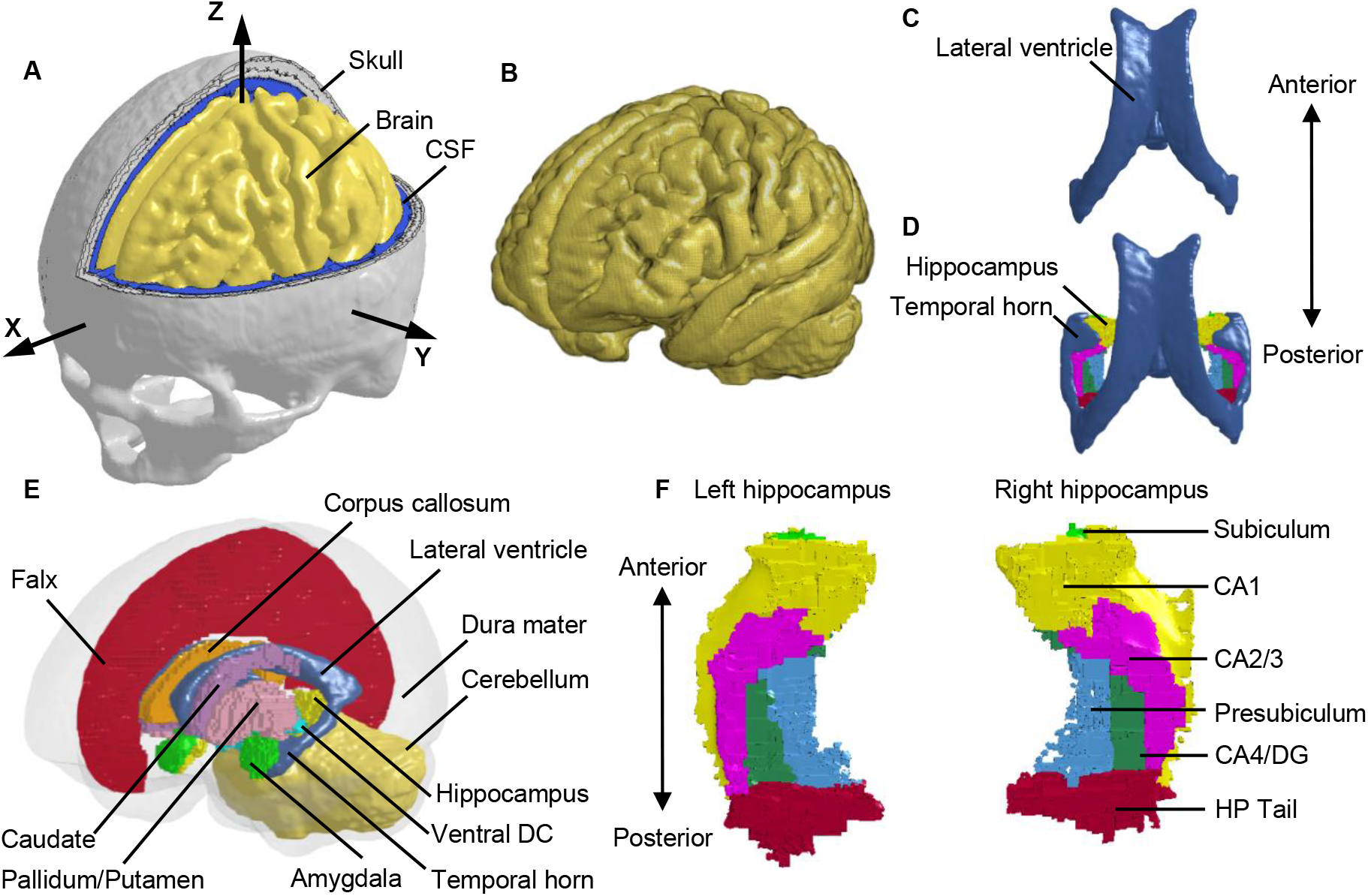
Finite element models of the human head with and without the temporal horn. (**A**) Head model with the skull open to expose the subarachnoid CSF and brain. A skull-fixed coordinate system and corresponding axes are illustrated with the origin at the center of gravity of the head. (**B**) Brain model with fine mesh. (**C**) Ventricles (i.e., lateral ventricles without the temporal horn, and third ventricle) in the NTH-model. (**D**) Ventricles (i.e., lateral ventricles with the temporal horn, and third ventricle) in the TH-model and hippocampus. (**E**) Isometric view of deep brain structures, cerebral ventricles, falx, and dura mater (in translucency) in the TH-Model. (**F**) Left and right hippocampal formations with subfields. CSF: cerebrospinal fluid; Ventral DC: ventral diencephalon; CA: cornu ammonis; DG: dentate gyrus; HP Tail: hippocampal tail.

To investigate the potential effect of the presence of the CSF-filled temporal horn on the hippocampus, we extended the NTH-model by adding the fluid-filled temporal horn to the cerebral ventricle (i.e., from Fig. 1C to Fig. 1D). This extended model (i.e., the TH-Model) has the same geometrical features, material properties, element formulation, and interface conditions as the NTH-Model, except for the newly added temporal horn. The volume ratio between the temporal horn and the brain in the TH-Model was 0.13%, falling within the range in healthy adults (0.1%-0.3%) (Bigler and Tate, 2001). Strain response and brain-skull relative motion estimated by the TH-Model were respectively evaluated by the experiments presented by Hardy et al. (2007) and Zhou et al. (2019c) in Appendix B. Details about the cerebral ventricle modeling and the brain-ventricle interface of the TH-Model are elaborated in the following two sections, along with that in the NTH-Model.

To facilitate the derivation of deformation-related metrics in regions of interest (ROIs) from completed simulations, the brain segmentation was registered to the coordinate system of the FE head model and then the brain elements were grouped into different sub-regions according to the spatial correspondence with the brain segmentation via an automated procedure implemented by a custom-built MATLAB script. For both the TH-Model and NTH-Model, the anatomically classified brain regions included cerebral cortex, cerebellum, hippocampus with six subfields as segmented by FreeSurfer 7 (i.e., cornu ammonis (CA) 1, CA2/3, CA4/dentate gyrus (DG), hippocampus tail (HP Tail), subiculum, and presubiculum) (Fig. 1F), and non-hippocampal paraventricular regions (i.e., amygdala, ventral diencephalon (ventral DC), pallidum, putamen, caudate, and corpus callosum (CC)) (Fig. 1E).

### Cerebral ventricle modeling

To emulate the fluid properties of the intraventricular CSF and potential CSF flow secondary to exterior loading, the cerebral ventricles in the TH-Model (Fig. 2A) and NTH-Model (Fig. 2B) were simulated using an ALE multi-material formulation. This formulation advances the solution in time using a two-step operation, in which the material is antecedently deformed in a Lagrangian step and subsequently followed by an advection step with the element variables being remapped (Zhou et al., 2019b). In the Lagrangian step, the intraventricular CSF deformation was determined by the equation of state (for dilatational responses) and constitutive equation (for deviatoric responses) listed in Table 1, together with associated formulations and material constants. In the advection step, a second-order van Leer scheme was selected, excelling in advection accuracy and numerical stability (van Leer, 1979).

**Table 1.** Material constant for the cerebral ventricles in the TH-Model and NTH-Model. *P* : pressure, *C* : intercept of *v*_*s*_ − *v*_*p*_ curves, *v s* : velocity of a shockwave traveling through the intermediary material, *v* _*p*_ : velocity of the shocked material; *S*_*1*_, *S*_*2*_, and *S*_*3*_: coefficients of the slope of the *v* _*s*_ − *v* _*p*_ curves, : Gruneisen gamma, *a* : first order volume correction to *γ*_0_; *ρ*_0_: initial density; *ρ* : instantaneous density;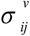 : deviatoric stress; *γ* : dynamic viscosity;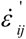: deviatoric strain rate; *PC*: cut-off pressure.

| Equation of state | $\rho_0$ (kg/m<sup>3</sup>) | $C$ (m/s) | $S_1$ | $S_2$ | $S_3$ | $a$ | $\gamma_0$ |
| --- | --- | --- | --- | --- | --- | --- | --- |
| $P = \frac{\rho_0 C^2 \mu (1 + (1 - \frac{\gamma^2}{2})\mu - \frac{a}{2} \mu^2)}{[1 - (S_1 - 1)\mu - S_2 \frac{\mu^2}{\mu + 1} - S_3 \frac{\mu^3}{(\mu + 1)^2}]}; \mu = \frac{\rho}{\rho_0} - 1$ | 1000 | 1482.9 | 2.10 | -0.17 | 0.01 | 0 | 1.2 |
| Constitutive equation | $\gamma$ (Pa.s) | $PC$ (MPa) | | | | | |
| $\sigma_{ij}^v = \gamma \dot{\epsilon}_{ij}$ | 0.001 | -22 | | | | | |

**Fig 2.**
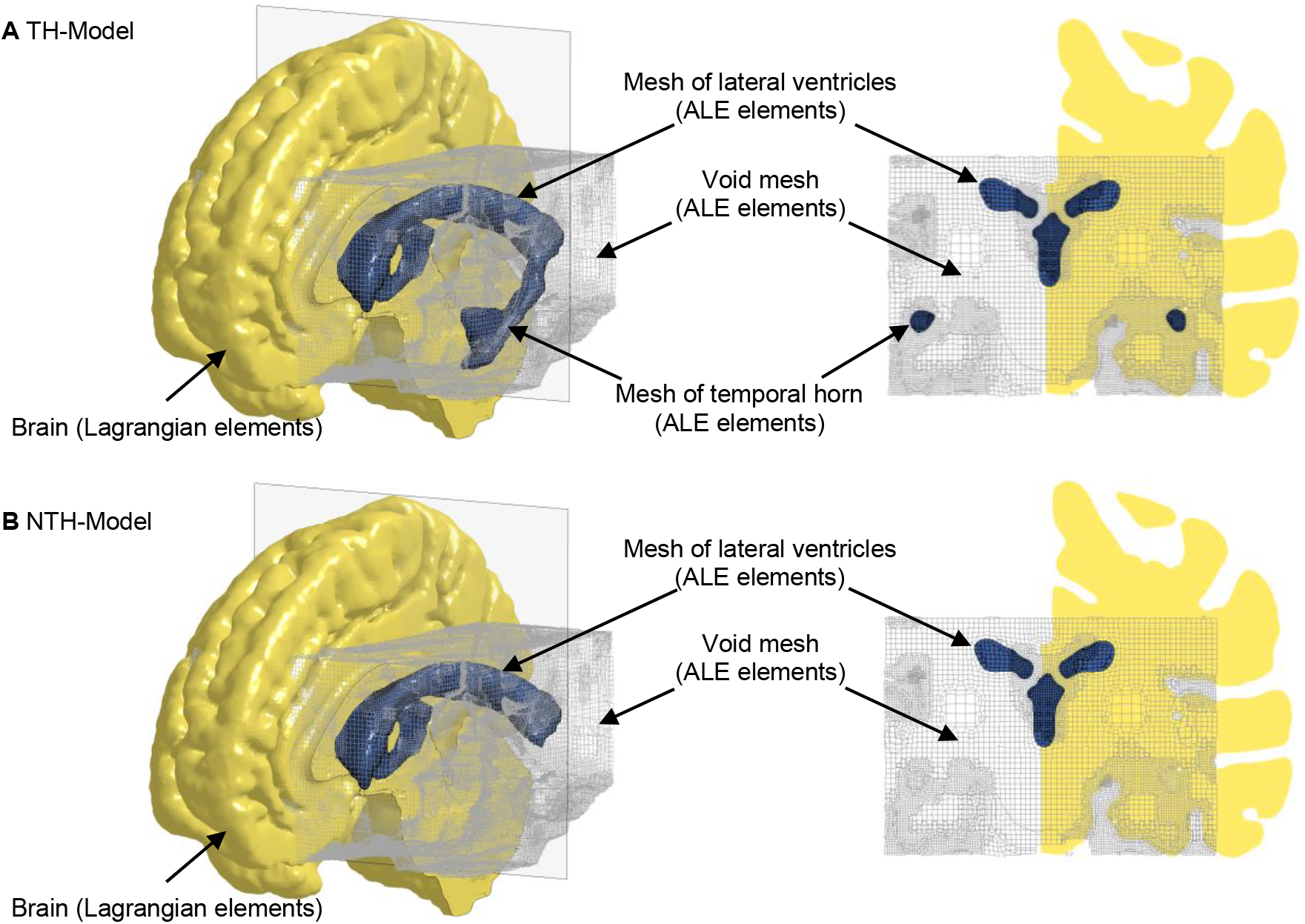
Brain-ventricle interfaces of the TH-Model (A) and NTH-Model (B). For each model, an isometric view of the brain model, the cerebral ventricle, and void mesh are shown on the left. Coronal sections at the planes indicated in the left subfigures are shown on the right. For better illustration, only half of the brain is visible. The cerebral ventricles are shown as blue shaded elements and the void mesh as wireframe elements. ALE: arbitrary Lagrangian-Eulerian.

### Brain-ventricle interface modeling

To couple the mechanical responses of the ALE-represented intraventricular CSF with the Lagrangian-represented brain, a penalty-based FSI coupling scheme (Batterbee et al., 2011;Zhou et al., 2019a) was implemented to both the TH-Model and NTH-Model. The implemented coupling scheme delivers tension and compression in the radial direction and allows relative motion in the tangential direction.

Owing to the requirement of implementing the penalty-based coupling scheme, any locations to which the fluid may potentially flow during the simulations are required to be meshed. Considering that the intraventricular CSF might flow to regions that were originally occupied by deep brain structures (due to deformation of the brain itself and the relative motion between the brain and cerebral ventricles during the simulation), additional meshes were generated in these regions, referred to as the “void mesh” in Fig. 2A and Fig. 2B, and initially overlapped with part of the brain elements. The void mesh was emulated with the ALE multi-material element approach, with material properties identical to that of the intraventricular CSF (Table 1) along with an extra void definition. Such a void definition ensured that no fluid was distributed within the void mesh under its initial configuration. The motion of the ALE elements followed the mass-weighted velocity in the ALE mesh (Hallquist, 2007).

### Loading conditions

Estimation of hippocampal response was obtained from the TH-Model and NTH-Model by simulating 6 representative football head impacts (Table 2 and Appendix B). At Stanford University, instrumented mouthguards have been developed to measure six-degree-of-freedom head kinematics during in-game head impacts to athletes (Liu et al., 2020;Cecchi et al., 2021). Using these instrumented mouthguards, over 500 head impacts in football have been video confirmed (Hernandez et al., 2015). In the current study, two concussive impacts, one with the athlete suffering alteration of consciousness (Case 1) and the other with the player having a milder but self-reported concussion (Case 2), and two sub-concussive impacts (Case 4 and Case 5) were simulated. In addition, a helmet-to-helmet collision involving two players was simulated with the struck player (Case 3) having a concussion and the striking player not (Case 6). Video recordings of the game were analyzed, through which the initial head kinematics were determined and further guided the laboratory reconstruction to obtain the dynamic kinematics of this collision (Pellman et al., 2003;Sanchez et al., 2019). All simulations were solved by the massively parallel processing version of LS-DYNA R11 double precision with 128 processors.

**Table 2.** Peaks of translational acceleration and rotational acceleration and injury severity of the 6 cases considered in this study. The X, Y, and Z axes are the same as those in the skull-fixed coordinate system in Fig. 1A. Note that Cases 1-2 and Cases 4-5 are on-filed impacts measured by the mouthguard (Hernandez et al., 2015), while Case 3 and Case 6 are laboratory-reconstructed impacts (Pellman et al., 2003;Sanchez et al., 2019).

| Case ID | Peak translational acceleration (g) |  |  |  | Peak rotational acceleration (krad/s<sup>2</sup>) |  |  |  | Injury severity |
| --- | --- | --- | --- | --- | --- | --- | --- | --- | --- |
|  | X | Y | Z | Magnitude | X | Y | Z | Magnitude |  |
| Case 1 | -40.6 | 100.4 | -63.4 | 106.1 | 12.89 | -3.06 | -3.24 | 12.95 | Concussion |
| Case 2 | -61.1 | -57.8 | -45.8 | 84.2 | 4.21 | 5.14 | -1.84 | 6.19 | Concussion |
| Case 3 | -31.9 | 133.4 | 41.6 | 134.0 | 4.65 | 1.20 | -6.81 | 7.50 | Concussion |
| Case 4 | -49.3 | -47.2 | -32.3 | 71.9 | 2.44 | -4.36 | -7.26 | 7.75 | Sub-concussion |
| Case 5 | 7.3 | 18.1 | 11.4 | 20.4 | 4.12 | 0.59 | 1.05 | 4.14 | Sub-concussion |
| Case 6 | -21.5 | -59.4 | 57.8 | 78.8 | -5.82 | -1.66 | -2.44 | 6.24 | Sub-concussion |

### Data analysis

For each computational simulation, the strains and strain rates in the 6 hippocampal subfields, the whole hippocampus, and 6 non-hippocampal periventricular regions were extracted from the TH-Model and NTH-Model, resulting in a total of 13 region-wise comparisons for each injury metric. This was motivated by the findings that hippocampal cell death was significantly affected by the strain (Cater et al., 2006) and hippocampal functional impairment was dependent on both strain and strain rate (Kang and Morrison, 2015) in *in vitro* TBI models on organotypic hippocampal slice cultures from rat. At each timestep, the element-wise strain and strain rate values were calculated as the first principal value of the Green-Lagrange strain tensor and the first principal value of rate of deformation tensor (Holzapfel, 2000). The element-wise strain and strain rate peaks were then identified as the maximum value of the strain and strain rate values across all timesteps. For each ROI, the element-wise strain and strain rate peaks of all affiliated elements were analyzed. To eliminate potential numerical artifacts (Panzer et al., 2012;Zhou et al., 2021b), the 95^th^ percentile values of element-wise strain peaks and element-wise strain rate peaks were respectively regarded as the strain peak and strain rate peak of the given region. A total of 13 ROIs, including 6 hippocampal subfields, the whole hippocampus, and 6 non-hippocampal paraventricular regions were considered in the current study. To quantify the variation in the responses per the inclusion of temporal horn, percentage differences in region-wise strain and strain rate peaks were computed for all ROIs in each loading case, with the value from the NTH-Model as reference. Similar postprocessing procedures have been implemented in previous studies (Gabler et al., 2018;Hajiaghamemar et al., 2020;Wu et al., 2021) to extract the regional-wise strain/strain rate peaks.

In total, 6 impacts were simulated by the TH-Model and NTH-Model, respectively. To statistically ascertain the influence of temporal horn on the deformation-related responses across the 6 impacts, the strain and strain rate peaks of all 6 impacts estimated by the TH-Model and NTH-Model were analyzed with a Wilcoxon matched-pairs signed-rank test (N=6), using an uncorrected significance threshold of p<0.05. This test was respectively implemented to all the 13 ROIs. Due to the small sample size, multiple comparisons correction was not performed in the current study.

## Results

### Strain and strain rate in the hippocampus and adjacent structures

We first aimed at elucidating the changes in strain and strain rate distribution due to the presence of the temporal horn. Cross-sections of whole-brain strain and strain rate maps are presented in Fig. 3. Almost identical strain and strain rate patterns were predicted by these two models, with the exception of strains over/approaching 0.2 (Fig. 3A-B) and strain rates over/approaching 30 s^-1^ (Fig. 3C-D)) around the temporal horn that was exclusively predicted by the TH-Model in all simulated loading cases.

**Fig. 3.**
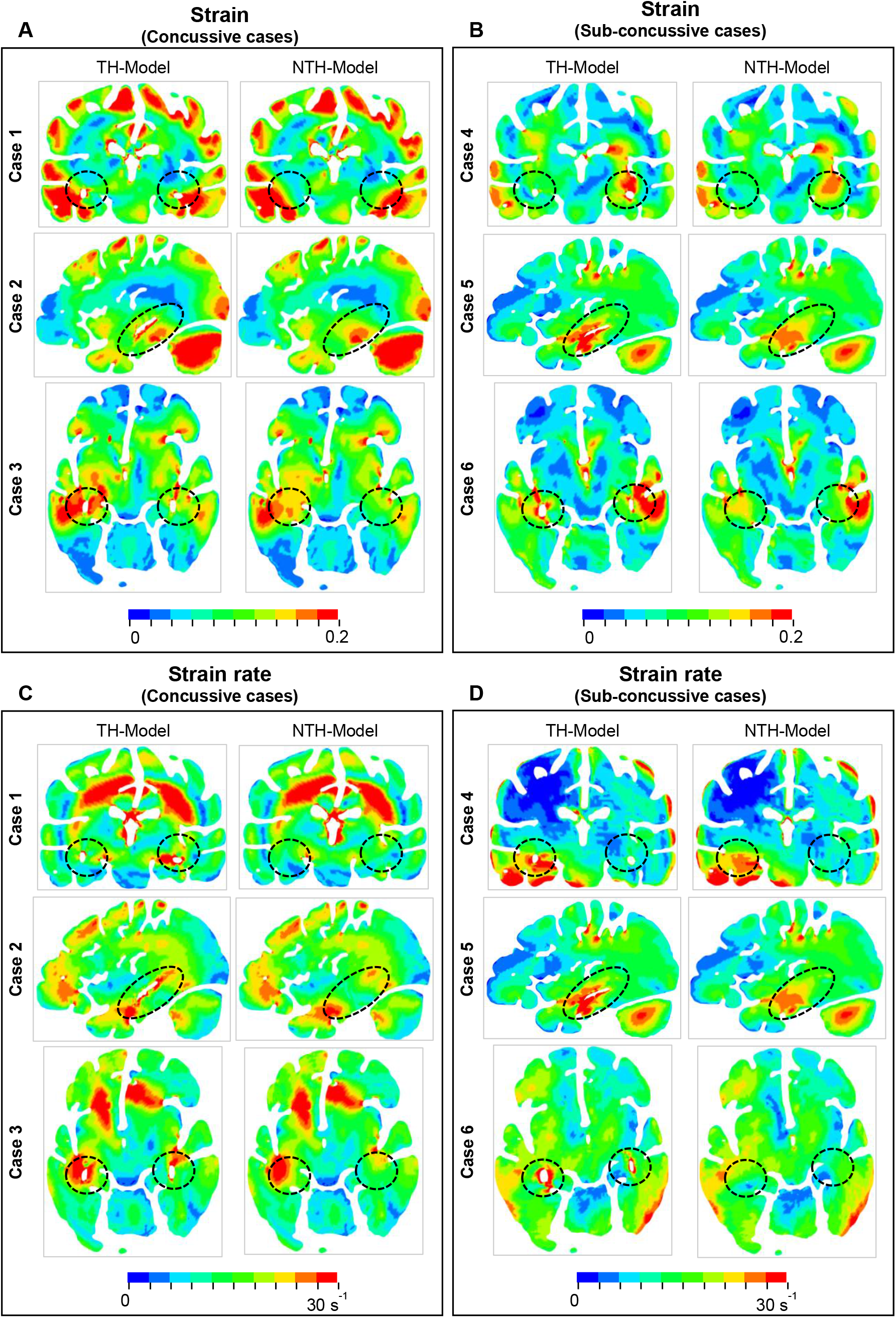
**Comparison of the maximum principal strain (A and B) and strain rate (C and D) distribution between the TH-model and NTH-model for three concussive and three sub-concussive impacts (Cases 1-3 and 4-6 respectively).** The temporal horn and adjacent tissue are highlighted by black dashed ellipses.

Close-up views of hippocampal strain and strain rate contours are presented in Fig. 4. Based on visual observation, regions experiencing strain approaching or over 0.2 in the TH-Model were more extensive with respect to their counterparts in the NTH-Model (Fig. 4A-B). This is particularly evident in CA1, CA2/3, and CA4/DG. Similarly, a more widespread distribution of strain rate approaching or over 30 s^-1^ was predicted by the TH-Model than the NTH-Model (Fig. 4C-D) in CA1, CA2/3, and HP Tail. This visual observation is quantitatively confirmed in Appendix D, in which larger volume ratios of strain over 0.2 and strain rate over 30 s^-1^ in the hippocampal subfields and the whole hippocampal level were predicted by the TH-Model with respect to the NTH-Model.

**Fig. 4.**
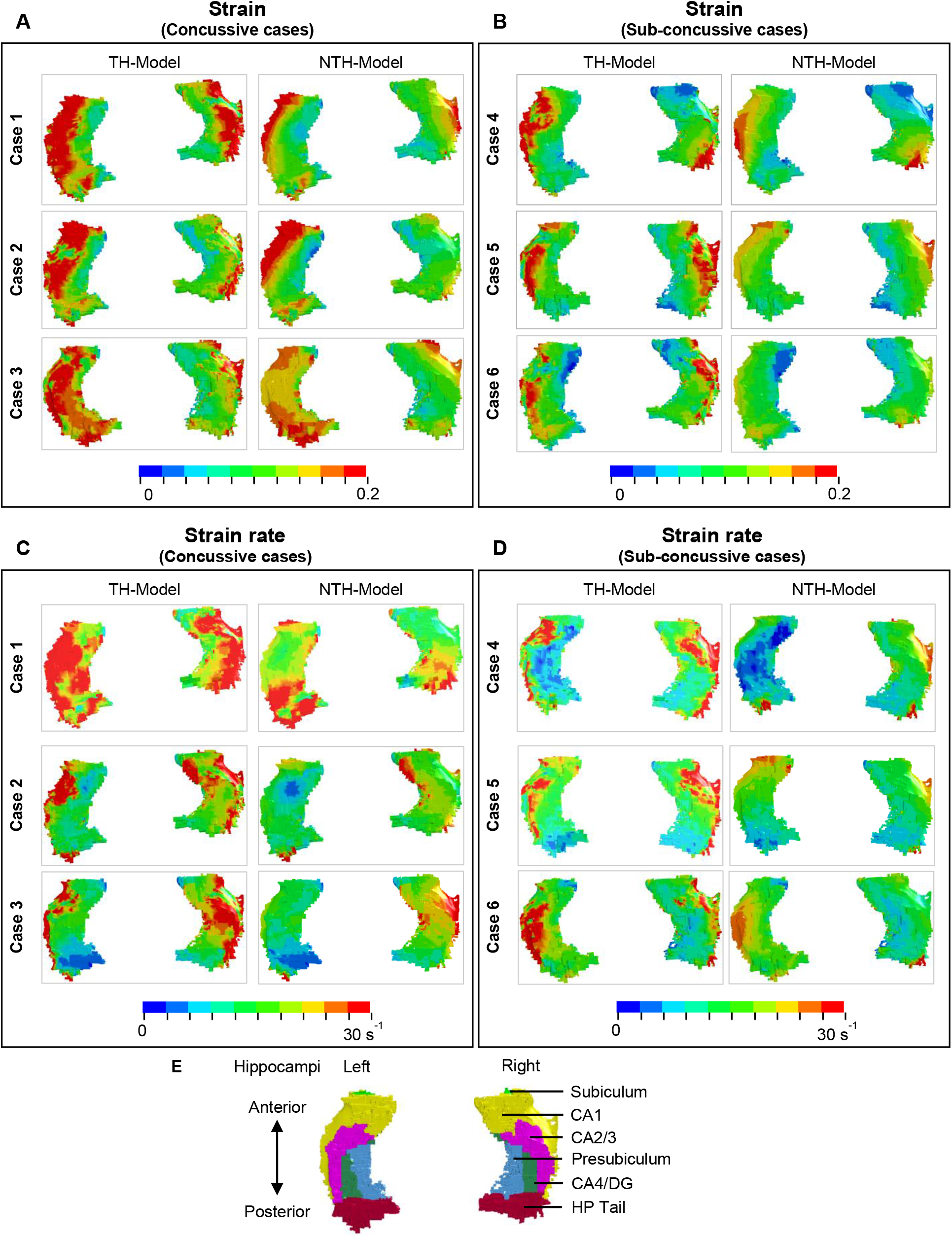
**Comparison of strain distribution (A and B) and strain rate distribution (C and D) in the hippocampi between the TH-model and NTH-model of three concussive impacts (Cases 1-3) and three sub-concussive impacts (Cases 4-6).** Subfigure (**E**) illustrates the hippocampal subfields. CA: cornu ammonis; DG: dentate gyrus; HP Tail: hippocampal tail.

Fig 5 shows a quantitative depiction of the findings in Fig. 4 with special focus on the peaking values: the addition of the temporal horn elevated the 95^th^ percentile maximum principal strain for almost all subfields and the whole hippocampus under all loading cases with the largest elevation (111.0 %) noted in CA2/3 in Case 5 (Fig. 5A-B). Similarly, the 95^th^ percentile maximum strain rate was increased per the addition of the temporal horn for almost all hippocampal subfields and the whole hippocampus, with the largest increase (168.0%) in HP Tail in Case 2 (Fig. 5C-D). Any decrements in strain or strain rate were less than 5%.

**Fig. 5.**
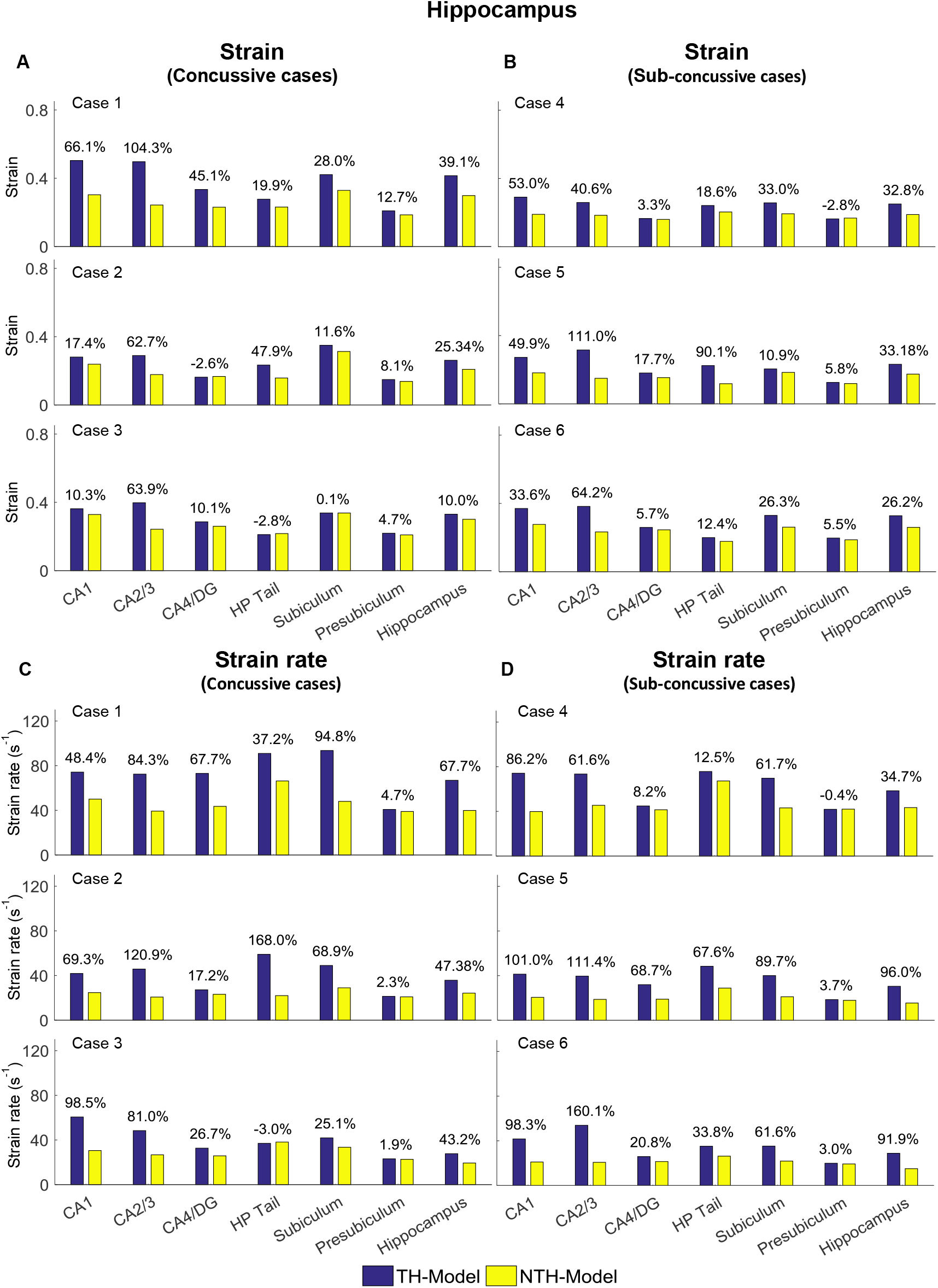
**Comparison of the 95_th_ percentile maximum principle strain and strain rate in the hippocampal subfields and the whole hippocampus between the TH-Model and NTH-model of 3 concussive impacts (Cases 1-3) and 3 sub-concussive impacts (Cases 4-6).**
(**A**) Comparison of strain in the hippocampal subfields of 3 concussive impacts. (**B**) Comparison of strain in the hippocampal subfields of 3 sub-concussive impacts. (**C**) Comparison of strain rate in the hippocampal subfields of 3 concussive impacts. (**D**) Comparison of strain rate in the hippocampal subfields of 3 sub-concussive impacts. Percentages in strain difference and strain rate difference are calculated with the results of the NTH-Model as the baseline. CA: cornu ammonis; DG: dentate gyrus; HP Tail: hippocampal tail.

We next aimed at identifying the anatomical regions most affected by the presence of the temporal horn. Using a Wilcoxon matched-pairs signed-rank tests on the region-wise strain and strain rate, we found considerable increases in strain (median value of percent strain difference>5%) on all six hippocampal subfields and the whole hippocampus at significant levels (p<0.05) (Table 3.A). For the strain rate, considerable increases (median value of percent strain rate difference>5%) were noted in all subfields except for the presubiculum.

**Table 3.** Wilcoxon matched-pairs signed-rank test on the region-wise strain and strain rate in the hippocampal subfields and whole hippocampus (A) and non-hippocampal regions (B) (N=6). Percentages in strain difference and strain rate difference between the TH-Model and NTH-model were calculated across all simulations and presented in the form of median and two quartile values with Q1 as 25^th^ percentile value and Q3 as 75^th^ percentile value. Note that N equals to the number of impacts simulated by each model. CA: cornu ammonis; DG: dentate gyrus; HP Tail: hippocampal tail; Ventral DC: ventral diencephalon; CC: corpus callosum.

| A | Regions | Percentage in strain difference |  | Percentage in strain rate difference |  |
| --- | --- | --- | --- | --- | --- |
|  |  | (median (Q1, Q3)) (%) | p | (median (Q1, Q3)) (%) | p |
|  | CA1 | 44.6 (33.6, 53.0) | 0.028 | 92.3 (69.3, 98.5) | 0.028 |
|  | CA2/3 | 64.6 (62.7, 104.3) | 0.028 | 97.9 (81.0, 121.0) | 0.028 |
|  | CA4/DG | 11.7 (3.2, 21.8) | 0.046 | 23.7 (17.2, 67.6) | 0.028 |
|  | HP Tail | 33.9 (18.6, 54.3) | 0.046 | 35.5 (12.5, 67.6) | 0.046 |
|  | Subiculum | 6.9 (5.5, 11.2) | 0.046 | 65.3 (61.6, 89.7) | 0.028 |
|  | Presubiculum | 19.9 (11.6, 28.0) | 0.028 | 2.7 (1.9, 3.7) | 0.046 |
|  | Hippocampus | 29.5 (25.3, 33.2) | 0.028 | 57.5 (34.7, 91.9) | 0.028 |
| B | Regions | Percentage in strain difference |  | Percentage in strain rate difference |  |
|  |  | (median (Q1, Q3)) (%) | p | (median (Q1, Q3)) (%) | p |
|  | Amygdala | 33.8 (17.1, 39.3) | 0.028 | 50.9 (40.4, 56.1) | 0.028 |
|  | Ventral DC | 8.2 (4.6, 12.2) | 0.028 | 9.35 (3.7, 13.1 ) | 0.028 |
|  | Pallidum | -1.7 (-4.2, 2.1) | 0.249 | -0.6 (-4.2, 4.4 ) | 0.753 |
|  | Putamen | -1.4 (-2.3, 2.8) | 0.249 | 2.2 (-0.4, 4.7) | 0.173 |
|  | Caudate | 1.5 (0.7, 5.5) | 0.917 | 0.1 (-2.5, 0.9) | 0.463 |
|  | CC | 0.7 (0.0, 1.3) | 0.249 | 2.5 (-0.4, 3.6) | 0.116 |

Among the non-hippocampal regions, both strain and strain rate were elevated in the TH-Model in the amygdala, which is along the anterosuperior border of the temporal horn, and to a lesser extent in the nearby ventral DC (Appendix E, Table 3.B). For the remaining more-distant regions, percentage differences in strain and strain rate were constantly less than 5% across the simulated loading cases.

### Stress in hippocampus and temporal horn

We then went on to explain the biomechanical reason for the hippocampal vulnerability. To ascertain the alteration of stress transmission associated with the temporal horn, Fig. 6A-B illustrates the maximum shear stress (i.e., a force triggering critical tissue deformation) endured by the temporal horn and hippocampus in the TH-Model and their counterparts in the NTH-Model, respectively. A much larger magnitude of shear stress in the hippocampus was noted in the TH-Model compared to the NTH-Model across all the cases (Fig. 6A). Conversely, the maximum shear stresses were less than 100 Pa in the temporal horn in the TH-Model, and over 1000 Pa in the temporal horn substitute in the NTH-Model (Fig. 6B). In addition, the distribution of shear stress within the hippocampus and temporal horn for one representative case (Case 2) are illustrated in Fig. 6C-D, in which a wider distribution of shear stress over 1000 Pa in the hippocampus was noted in the TH-Model compared to the NTH-Model. It is thus indicated that an altered stress transmission associated with the temporal horn causes elevations in strain and strain rate.

**Fig. 6.**
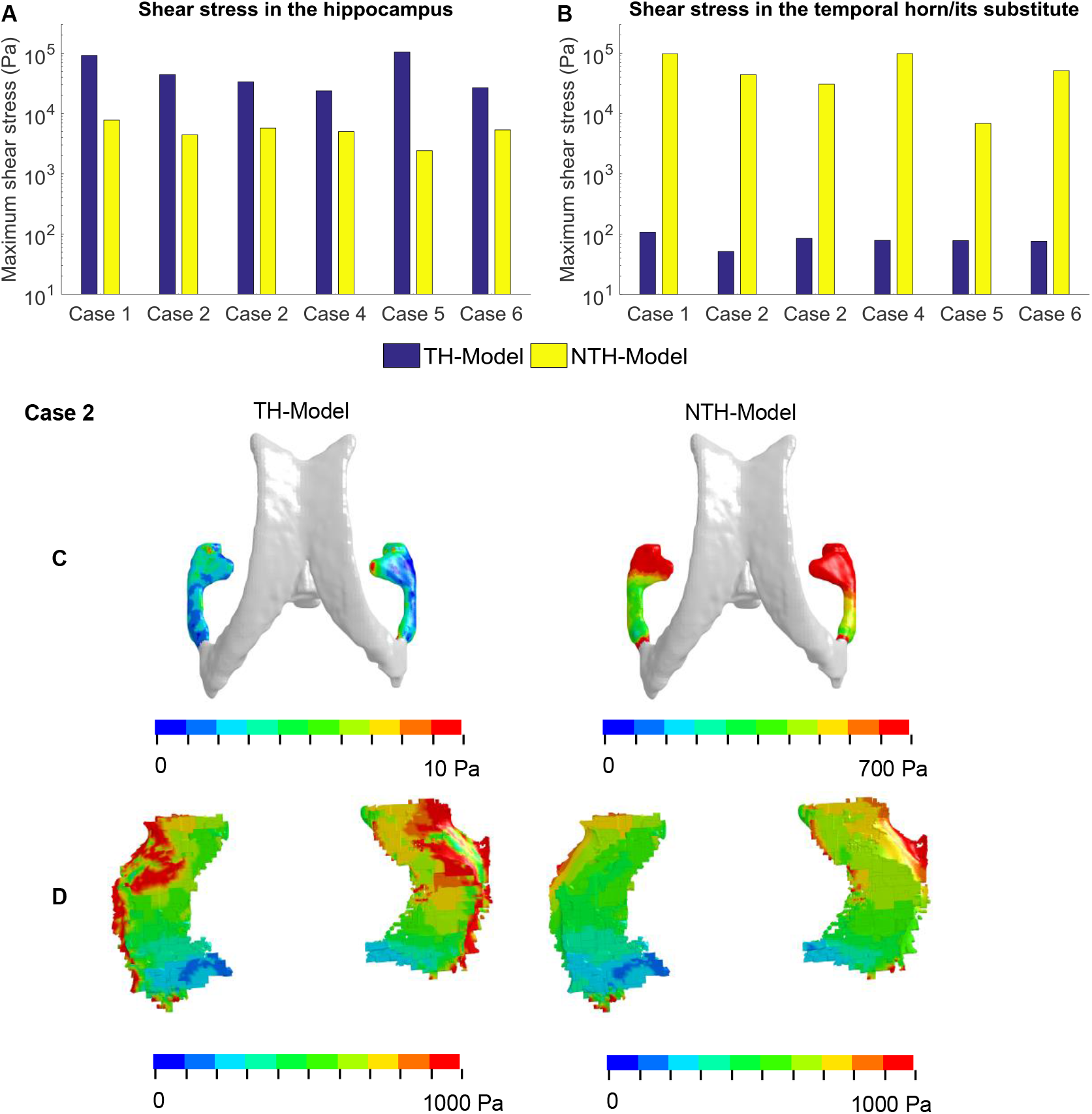
**Maximum shear stresses in the hippocampus (A) and temporal horn/its substitute (B) predicted by the TH-Model and NTH-Model in 6 cases; (C) Contours of maximum shear stress in the CSF within the temporal horn in the TH-Model and its substitute in the NTH-Model; (D) Contours of maximum shear stress endured by the hippocampi in the TH-Model and NTH-Model.** Note that, in the NTH-Model, the temporal horn is modeled as brain, not fluid.

## Discussion

The current study attempted to elucidate why the hippocampus is so commonly affected by brain trauma. We used two FE models: one with and the other without the temporal horn, and incorporated an anatomically accurate description of temporal horn, a mechanically realistic representation of intraventricular CSF as fluid elements, and a fluid-structure interaction coupling approach for the brain-ventricle interface. The presence of the temporal horn not only extended the distribution of high strains and strain rates in the surrounding area, but also increased their magnitude in the hippocampus, particularly in the subfields of CA1, CA2/3, HP Tail, subiculum, and presubiculum. Other adjacent regions including the amygdala and ventral DC showed similarly increased strain and strain rate with the presence of the temporal horn, but distant regions (e.g., corpus callosum) did not. These computational findings suggest that the presence of the temporal horn likely exacerbates the biomechanical vulnerability of the hippocampus following head impacts.

This biomechanical finding correlates well with the prevalence of hippocampal trauma in humans data and animal biomechanical models. Several postmortem neuropathological studies (Kotapka et al., 1992;Kotapka et al., 1993;Kotapka et al., 1994;Maxwell et al., 2003) have detected overt neuronal damage/loss in the hippocampus of TBI victims with high incidence rates up to 73%-87% (although the exact loadings endured were lacking). Animal models employing custom-built pneumatic devices that deliver impulsive angular accelerations, similar to the loading mode in the current study, have shown hippocampal lesions in non-human primates (Gennarelli et al., 1982;Kotapka et al., 1991), which have a similar hippocampal morphology and spatial relationship to the temporal horn (Insausti and Amaral, 2003;Amaral et al., 2007). A version of the device modified to deliver impulsive loading caused selective hippocampal damage to porcine brains (Smith et al., 1997) (which again have a similar relationship between the hippocampus and temporal horn to humans (Félix et al., 1999)). Thus, animal models with similar morphological relationships between the temporal horn and hippocampus support a biomechanical link between the two.

Our computational results predicted an altered stress transmission associated with the temporal horn, providing an explanation for the elevations in strain and strain rate in the TH-Model. As illustrated in Fig. 6B, the shear stress endured by the temporal horn in the TH-Model was less than 100 Pa, which realistically reflected the low shear resistance nature of CSF. Comparatively, the shear stress experienced by the substitute of the temporal horn (i.e., brain parenchyma) in the NTH-Model was over 1000 Pa, providing an unrealistic interaction with the neighboring tissue. These regions adjacent to the temporal horn (such as the hippocampus, amygdala, and ventral DC, as are the ROIs in the current study) were easier to deform when associated with the addition of the temporal horn in the TH-Model, consequently exacerbating the strain and strain rate in these ROIs. This explanation was further verified in Fig. 6A, where the shear stress endured by the hippocampus was larger in the TH-Model, consistent with an amplified force exerted on the hippocampus with the addition of the temporal horn.

Two previous computational studies simulated football head impacts, consistently reporting an increased susceptibility of the hippocampus to injury (Viano et al., 2005;Zhao et al., 2017). However, the ventricular elements in these two models and other ones (Kleiven, 2007;Mao et al., 2013;Atsumi et al., 2018;Trotta et al., 2020) were manually picked with reference to the brain atlas, lacking mesh conformity of the anatomic ventricle profile. Our work used a novel FE model of the brain that involves orders of magnitude more elements than used in typical models (e.g., millions instead of thousands), enabling a realistic depiction of the geometrical features of the temporal horn. Intraventricular CSF elements in existing head models (Viano et al., 2005;Kimpara et al., 2006;Takhounts et al., 2008;Mao et al., 2013;Ho et al., 2017;Zhao et al., 2017;Zhou et al., 2019a;Li et al., 2020) are predominantly represented by Lagrangian elements, with the mesh following the material deformation without material advection, neglecting the potential fluid flow during the impact. Here, we leveraged a fluid element formulation (i.e., ALE multi-material formulation) for the cerebral ventricle, emulating the fluid properties of intraventricular CSF and potential fluid flow following external stimuli. To couple the mechanical responses of the ALE-represented ventricular CSF elements with the Lagrangian-represented brain elements, a penalty-based coupling was implemented. Such a coupling algorithm permits relative motion in the tangential direction and delivers tension and compression in the radial direction, circumventing severe element distortion at the interfacial boundary. The FSI approach excels in not only realistically representing the fluid behavior of the CSF but also maintaining numerical stability without causing severe element distortion, supporting its validity for the current application. Nevertheless, it is worth clarifying that our data suggest that omitting the temporal horn, as is the case in most existing head models, may still be acceptable for these studies that focus on regions far from the temporal horn (e.g., corpus callosum, caudate, putamen, pallidum).

Hippocampal cell death tolerance criteria were presented by Cater et al. (2006) by relating three independent variables (i.e., strain in the range of 0.05 to 0.5, hippocampal subfields, time post-injury) to resultant cell death under *in vitro* conditions via mathematical equations, which were valid within the strain rate regime of 0.1-50 s^-1^. Similarly, another *in vitro* study reported tolerance criteria for hippocampal function impairment in the form of mathematical formulations between input mechanical stimuli (i.e., strain up to 0.44 and strain rate up to 30 s^-1^) and output electrophysiological alterations (Kang and Morrison, 2015). In the current study, the hippocampal responses predicted by the TH-Model peaking from 0.29 to 0.50 for strain and from 53.9 s^-1^ to 93.6 s^-1^ for strain rate in the six simulated impacts. The range of FE-derived strains and strain rates reached the criteria of electrophysiological impairment and cell death as aforementioned.. However, it should be noted that certain disparities existed between the data ranges of the current computational results and the loading regimes from which these two hippocampus-related tolerance criteria were fitted. Moreover, the cultured hippocampal slices in Cater et al. (2006) and Kang and Morrison (2015) were obtained from the rat brain. Extrapolation of the tolerance criteria derived from the animal brain under *in vitro* conditions to the human brain under *in vivo* conditions requires further verification (Seok et al., 2013).

While the presence of the CSF-filled temporal horn may be a contributing factor for the hippocampal vulnerability, additional mechanisms, such as the selective vulnerability of hippocampal neurons to hypoxemia and ischemia (Pulsinelli, 1985;Ng et al., 1989) and pathological neuronal excitation involving glutamate and other excitatory amino acid neurotransmitters (Faden et al., 1989;Bullock et al., 1990), may play important roles in human hippocampal injury. We suggest that the adverse effects of the temporal horn during the primary impact, the superimposed hypoxia/ischemia and neuroexcitotoxicity secondary to the impact, as well as other potential unknown mechanisms, collectively contribute to the hippocampal vulnerability.

### Limitations and future work

Although the current study yielded some new insights into the biomechanical dependency of the hippocampus on the temporal horn, certain limitations exist which require further investigation. First, only 6 representative sports-related inertial impacts were simulated in the current study with the severities at concussive and sub-concussive levels. A systematic investigation that covers more impact-related variables (e.g., impact duration, impact directions, rotational velocity) with their magnitudes spanning over the regimes measured from the realistic impacts is planned for future work to identify the critical scenarios that the temporal horn exhibited a more pronounced effect on the hippocampus. Moreover, caution should be exercised when extrapolating the current findings obtained from concussive and sub-concussive impacts to extra injury scenarios (e.g., fatal brain injury, penetrating head injury).

Secondly, due to the computational challenges, the brain-skull interfaces in both models in the current study were simulated by approximating the subarachnoid CSF as a Lagrangian-represented structure. Given that the ROIs in the current study are all located at central brain regions, the influence exerted by the brain-skull influence on the deep brain structures was expected to be limited (Kleiven and Hardy, 2002). Per the benefits of using ALE elements for the cerebral ventricles, the impact-induced fluid flow was considered, but not quantified in the current study. A detailed examination of flow patterns of CSF remains to be appropriately quantified in the future (Lang and Wu, 2021).

Thirdly, to incorporate explicit representations of the hippocampal subfields in the FE models, Freesurfer was used to segment the MRI with a resolution of 1 × 1× 1 mm^3^ to take advantage of the isotropic high-resolution atlas and incorporate this detailed isotropic segmentation into the FE model. Such a software choice was for the consistency purpose, since the brain profile used for the development of FE model was obtained from Freesurfer. However, it should be highlighted there are many different segmentation methods for hippocampal subfields, presenting certain variances in specific subfield delineation (Yushkevich et al., 2015;Wisse et al., 2021). Thus, caution should be exercised when using Freesurfer for hippocampal subfield segmentation (Wisse et al., 2014). In fact, there appear no approaches with guaranteed utility and validity to segment hippocampal subfield from isotropic 1 mm^3^ MRI (Wisse et al., 2021), as is the case for the subfield delineation in the FE model. This segment-induced error inevitably compromised the accuracy of hippocampal subfield representations in the FE model, which is a limitation of the current study. Nevertheless, compared with the studies in which the hippocampus was treated as a single medium (Takhounts et al., 2008;Mao et al., 2013;Miller et al., 2016;Atsumi et al., 2018;Li et al., 2020;Trotta et al., 2020;Zhou et al., 2020a), the current work made the first step to differentiate the hippocampal substructures in the FE model of the human brain. Future work is planned to further refine the model towards an anatomically more authentic hippocampal subfield representation.

Lastly, due to the lack of neuroimaging data of these players with their head impacts being simulated, it is hardly possible to ascertain whether hippocampal injury indeed occurred in these six simulated impacts. At Stanford, ongoing effort is dedicated to deploy instrumented mouthguards to football players, obtaining real-time measurements of the impacts sustained by these players (Camarillo et al., 2013;Hernandez et al., 2015;Domel et al., 2020;Liu et al., 2020). This information is complemented by medical imaging of the football players pre- and post-impact (Parivash et al., 2019;Mills et al., 2020). Findings in the current work will be further testified by correlating on-field football impacts, to computationally predicted hippocampal deformation, to image-based evidence of hippocampal injury.

## Conclusion

This study investigated the biomechanical mechanism of hippocampal injury associated with the presence of the temporal horn by leveraging two models, with and without the inclusion of the temporal horn. The results showed that the temporal horn has a significant biomechanical effect in the surrounding area and induces increased magnitudes of the strain and strain rate in the hippocampus throughout its subfields, identifying the temporal horn as a contributing factor to the hippocampal vulnerability. This study suggests that proper modeling of the temporal horn be considered when developing mechanical tolerance and designing protective strategies specifically for the hippocampus.

## Acknowledgements

Drs. Michael Zeineh, Gerald Grant, and David Camarillo received funding from the Pac-12 Conference’s Student-Athlete Health and Well-Being Initiative and Taube Stanford Children’s Concussion Initiative. Drs. Svein Kleiven and Xiaogai Li received funding from the Swedish Research Council (VR-2016-05314 and VR-2016-04203), while Dr. Marios Georgiadis received funding from the Swiss National Science Foundation (P400PM_180773). Zhou Zhou received funding from KTH Royal Institute of Technology (Stockholm, Sweden). The content of this article is solely the responsibility of the authors and does not necessarily represent the official views of funding agencies. The simulations were performed on resources provided by the Swedish National Infrastructure for Computing (SNIC) at the center for High Performance Computing (PDC). Gong Jing at the PDC center is acknowledged for the technical maintenance of needed software employed in the current study. In addition, the authors thank Dr. Erin Bigler at the University of Utah School of Medicine for helpful discussion and the reviewers for the stimulating comments and valuable suggestions that substantially improved this paper.

## Appendix A Development of finite element head model without the temporal horn

The finite element (FE) head model without the temporal horn (i.e., the no-temporal-horn (NTH)-Model) used in this study was previously established at KTH Royal Institute of Technology in Stockholm, Sweden (Zhou et al., 2020a). The geometry of the head model was extracted from an averaged magnetic resonance imaging (MRI) head template database (Fillmore et al., 2015). High-resolution T1- and T2-weighted images were segmented using the Freesurfer 7 (Fischl, 2012). The segmentation was subsequently processed by the 3D Slicer (Pieper et al., 2004) to obtain the surfaces of the skull, the brain, the third and the lateral ventricles, with the temporal horn being disregarded. All surfaces then served as an input to the Hexotic software, generating all hexahedron elements based on an octree algorithm (Maréchal, 2009). The falx and tentorium, which were almost invisible in the MRI, were manually created as shell elements based on the anatomical illustrations, while the pia mater and dura mater were generated by finding the outer faces of brain elements and subarachnoid cerebrospinal fluid (CSF) elements, respectively.

The material representation of simulated head components is summarized in Table A1 and Table A2. For the brain, a second-order Ogden-based hyperelastic constitutive law was used to describe the nonlinear behavior of the brain tissue, with additional linear viscoelastic terms to account for rate dependence. The subarachnoid CSF was modeled as a nearly incompressible material and shared interfacial nodes with the brain and skull. Mechanical properties of the intracranial membrane (i.e., pia mater and falx/tentorium/dura mater) were determined from the averaged material stress-strain curves from the tissue experiments. The brain, subarachnoid CSF, and intracranial membranes were modeled as Lagrangian elements (Zhou et al., 2020a). In particular, the ventricles were represented by arbitrary Lagrange-Eulerian (ALE) fluid elements, and their responses were coupled to the brain via a fluid-structure interaction (FSI) scheme, which is detailed in the “Cerebral ventricle modeling” and “Brain-ventricle interface modeling” sections of the current study. Note that, in the NTH-model, the temporal horn was substituted as brain parenchyma with the material constants in Table A2.

**Table A1.** Material properties for the finite element head model. K is Bulk modulus and N/A is not applicable.

| Tissue | Young's modulus (MPa) | Density (kg/dm3) | Poisson's ratio | Reference |
| --- | --- | --- | --- | --- |
| Cortical bone | 15000 | 2.00 | 0.22 | (Kleiven, 2007) |
| Porous bone | 1000 | 1.3 | 0.24 | (Kleiven, 2007) |
| Brain | Hyper-Viscoelastic (Table A2) | 1.04 | 0.5 | (Kleiven, 2007) |
| CSF/Ventricle | K = 2.1 GPa | 1.00 | N/A | (Kleiven, 2007) |
| Dura/Falx/Tentorium | Average stress-strain curve | 1.13 | N/A | (Aimedieu and Grebe, 2004) |
| Pia | Average stress-strain curve | 1.13 | N/A | (Van Noort et al., 1981) |

**Table A2.** Ogden hyperelastic and liner viscoelastic constants for the brain material modeling. μ_i_ and α_i_ are Ogden parameters, G_i_ represents the 6 shear relaxation moduli, β_i_ are the 6 decay constants.

| Parameter | Value | Parameter | Value |
| --- | --- | --- | --- |
| $\mu_1$ (Pa) | 53.8 | $\alpha_1$ | 10.1 |
| $\mu_2$ (Pa) | -120.4 | $\alpha_2$ | -12.9 |
| $G_1$ (MPa) | 0.32 | $\beta_1$ (s<sup>-1</sup>) | 10<sup>6</sup> |
| $G_2$ (kPa) | 78 | $\beta_2$ (s<sup>-1</sup>) | 10<sup>5</sup> |
| $G_3$ (kPa) | 6.2 | $\beta_3$ (s<sup>-1</sup>) | 10<sup>4</sup> |
| $G_4$ (kPa) | 8.0 | $\beta_4$ (s<sup>-1</sup>) | 10<sup>3</sup> |
| $G_5$ (kPa) | 1.0 | $\beta_5$ (s<sup>-1</sup>) | 10<sup>2</sup> |
| $G_6$ (kPa) | 3.0 | $\beta_6$ (s<sup>-1</sup>) | 10<sup>1</sup> |

## Appendix B Validation of brain-skull relative motion and brain strain

The strain response and brain-skull relative motion estimated by the model with the temporal horn (i.e., the TH-Model) was validated against the available experimental data presented by Hardy et al. (2007) and Zhou et al. (2019c). In Hardy et al. (2007), a high-speed, biplane X-ray system was used to track the motion of the radiopaque neutral density targets (NDTs) implanted in cluster array within the cadaveric brain. The NDT initial coordinates and motion were obtained with respect to an anatomical coordinate system with the c.g. of the head being the origin. Strain in the volume encompassed by the NDT cluster was calculated by imposing the experimentally measured NDT motions to an auxiliary model that was developed by connecting each NDT to its neighboring counterparts to form tetra elements (Zhou et al., 2019c).

In the present work, three representative cases are selected, including C288-T3 (sagittal impact), C380-T1 (coronal impact), and C380-T2 (horizontal impact). To numerically reproduce the experimental impacts, the recorded head kinematic curves were imposed to the node which locates at the center of gravity of the corresponding cadaveric head and is rigidly attached to the skull. To approximate the specimen anthropometry, the model was scaled independently in directions of both the depth and breadth to match the reported cadaveric head sizes. The node nearest to the start position of an experimental NDT target was taken as the marker location in the model. Motions of the identified nodes with respect to the skull along three anatomical coordinate directions are obtained from the whole head model simulation with the detailed results presented in **Fig. B1-B3**. Following the procedures established by Zhou et al. (2019c), the initial positions of the identified nodes and the nodal motion responses predicted in the model was used to calculate the strain responses, specifically first principal Green-Lagrange strain and shear Green-Lagrange strain, of the brain model. The strain validation results are shown in **Fig. B4**.

**Fig. B1.**
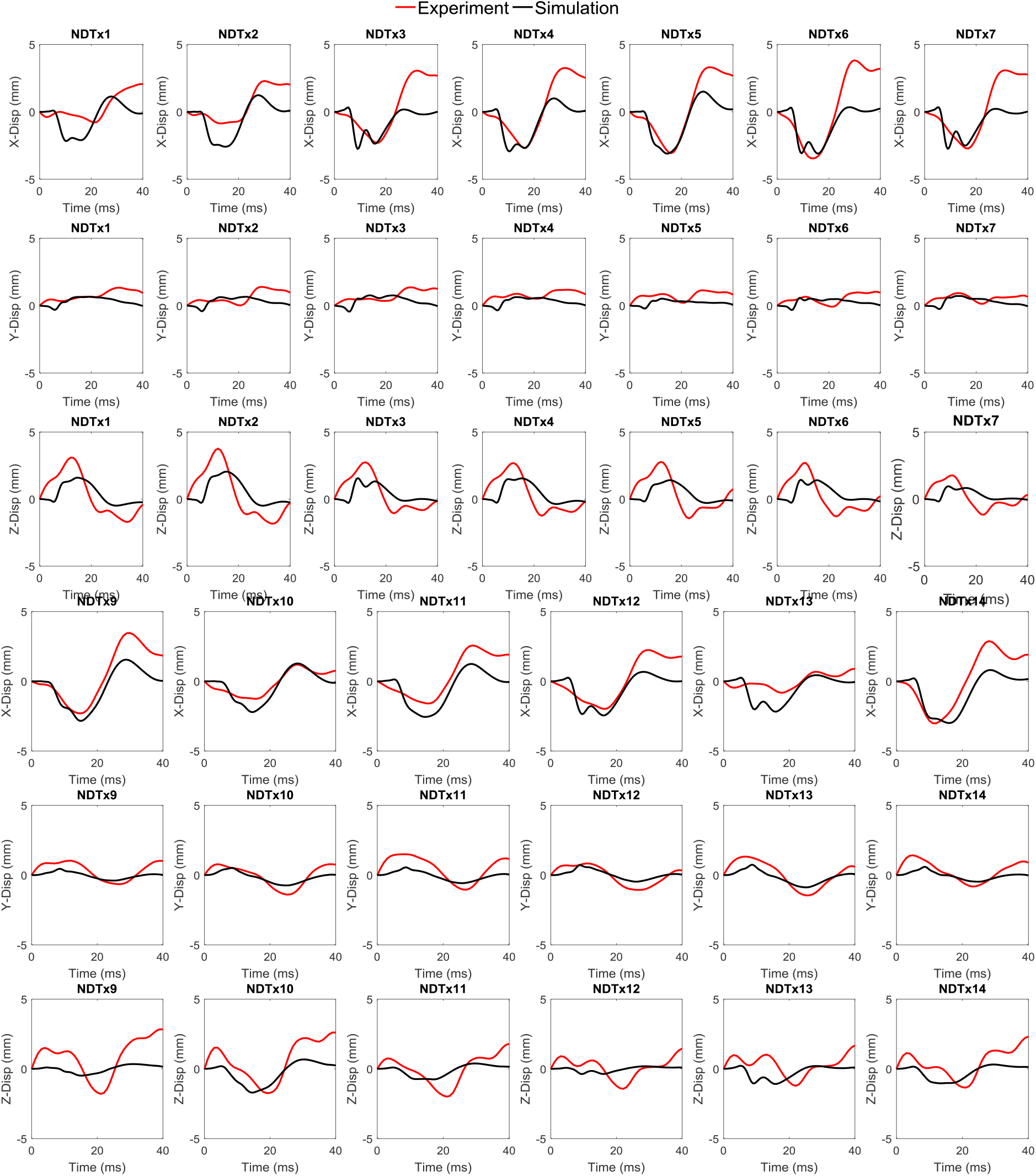
Comparison between experimental and simulated brain-skull relative motion for the experiment C288-T3.

**Fig B1.**
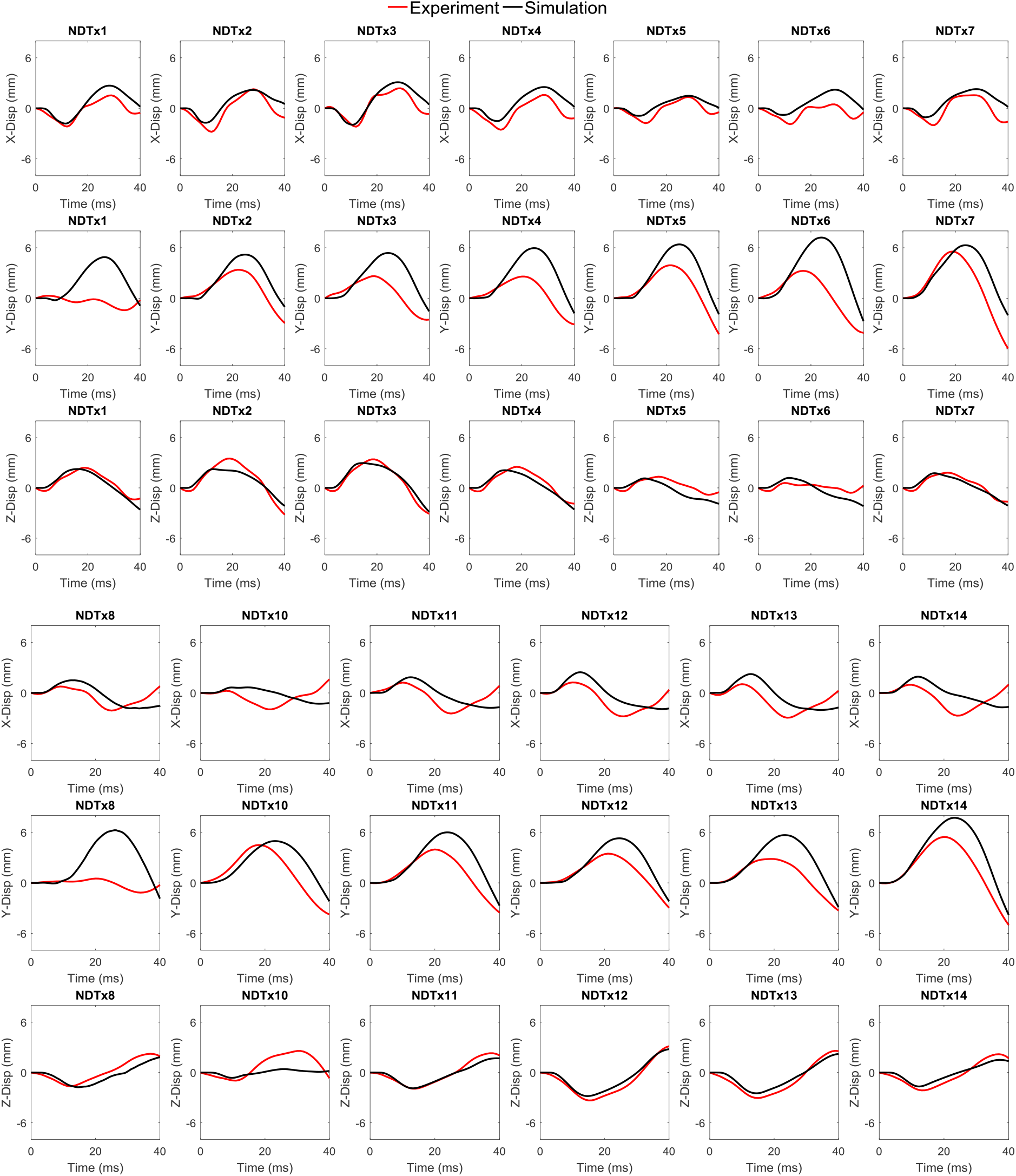
Comparison between experimental and simulated brain-skull relative motion for the experiment C380-T1.

**Fig B3.**
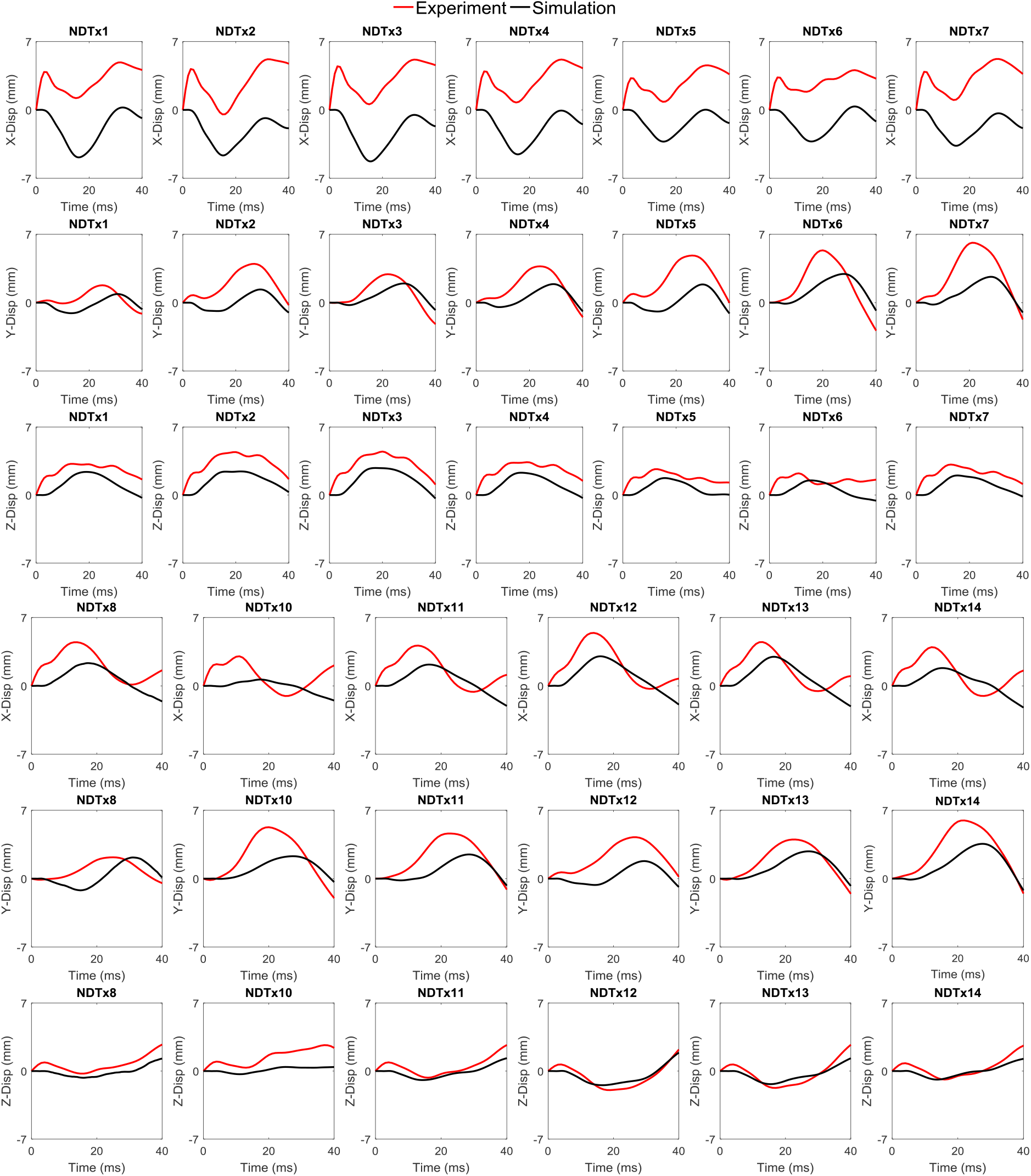
Comparison between experimental and simulated brain-skull relative motion for the experiment C380-T2.

**Fig B4.**
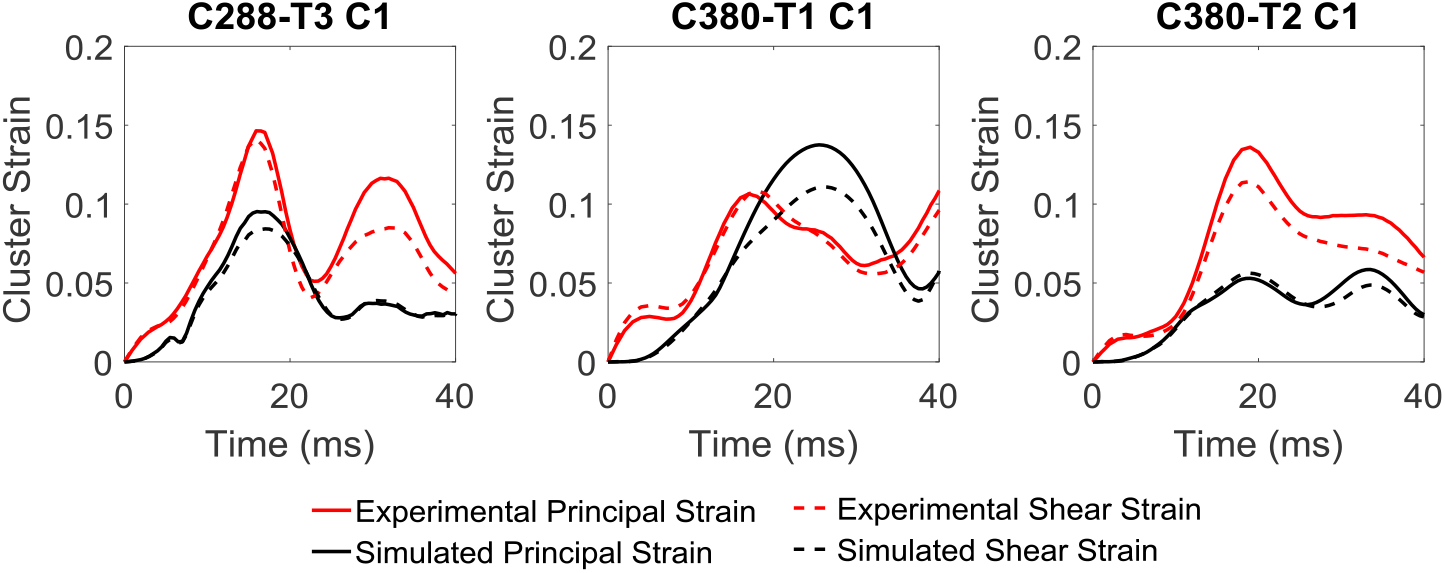
Comparison between experimental and simulated brain strains.

## Appendix C Loading curves for 3 concussive impacts and 3 sub-concussive impacts

**Fig C1.**
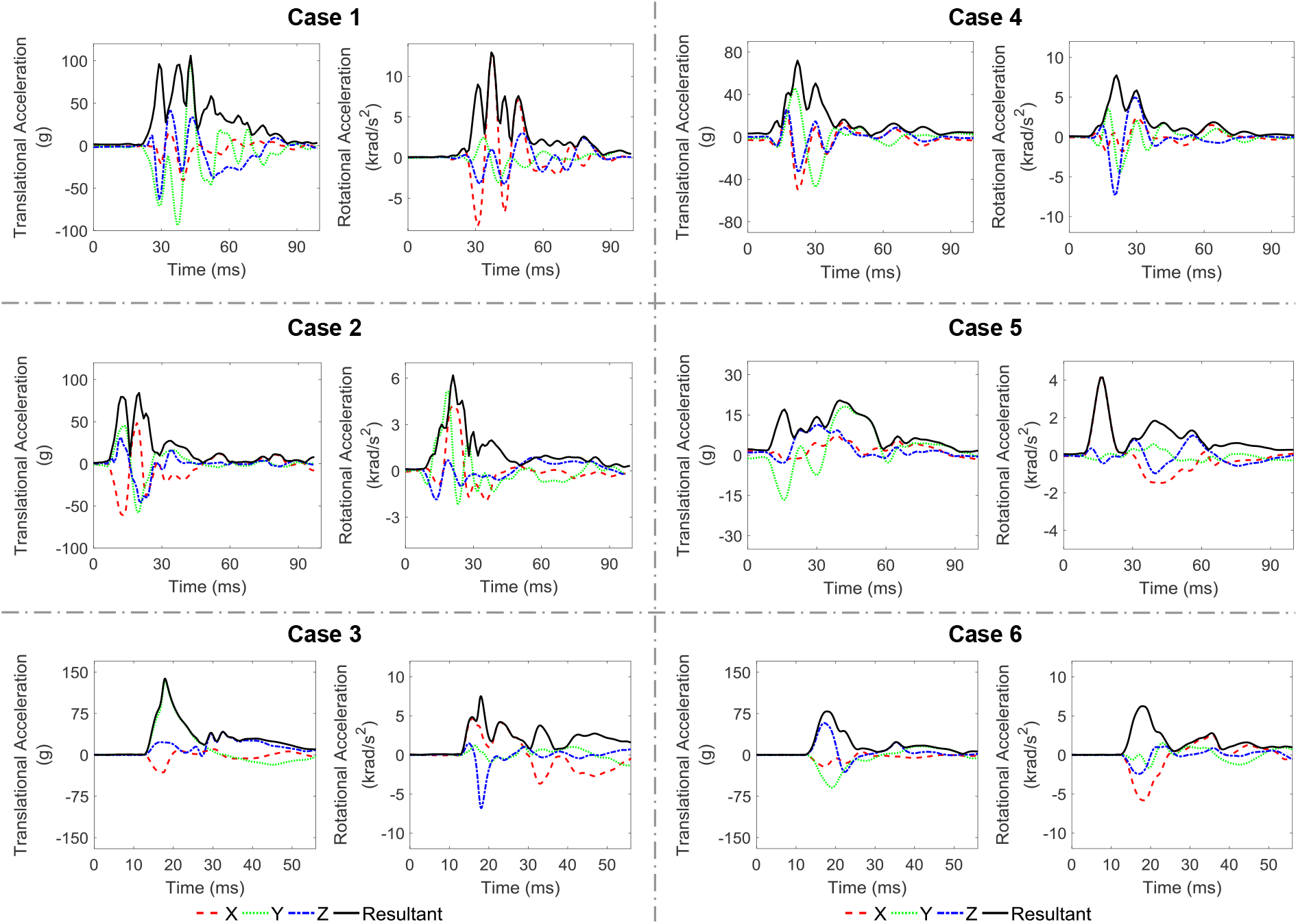
Loading conditions for 3 concussive impacts (Cases 1-3) and 3 sub-concussive impacts (Cases 4-6). The X, Y, and Z axes are the same as those in the skull-fixed coordinate system in Fig. 1a.

## Appendix D Volume ratio of the hippocampal subfields and whole hippocampus with strain over 0.2 and strain rate over 30 s^-1^

**Fig D1.**
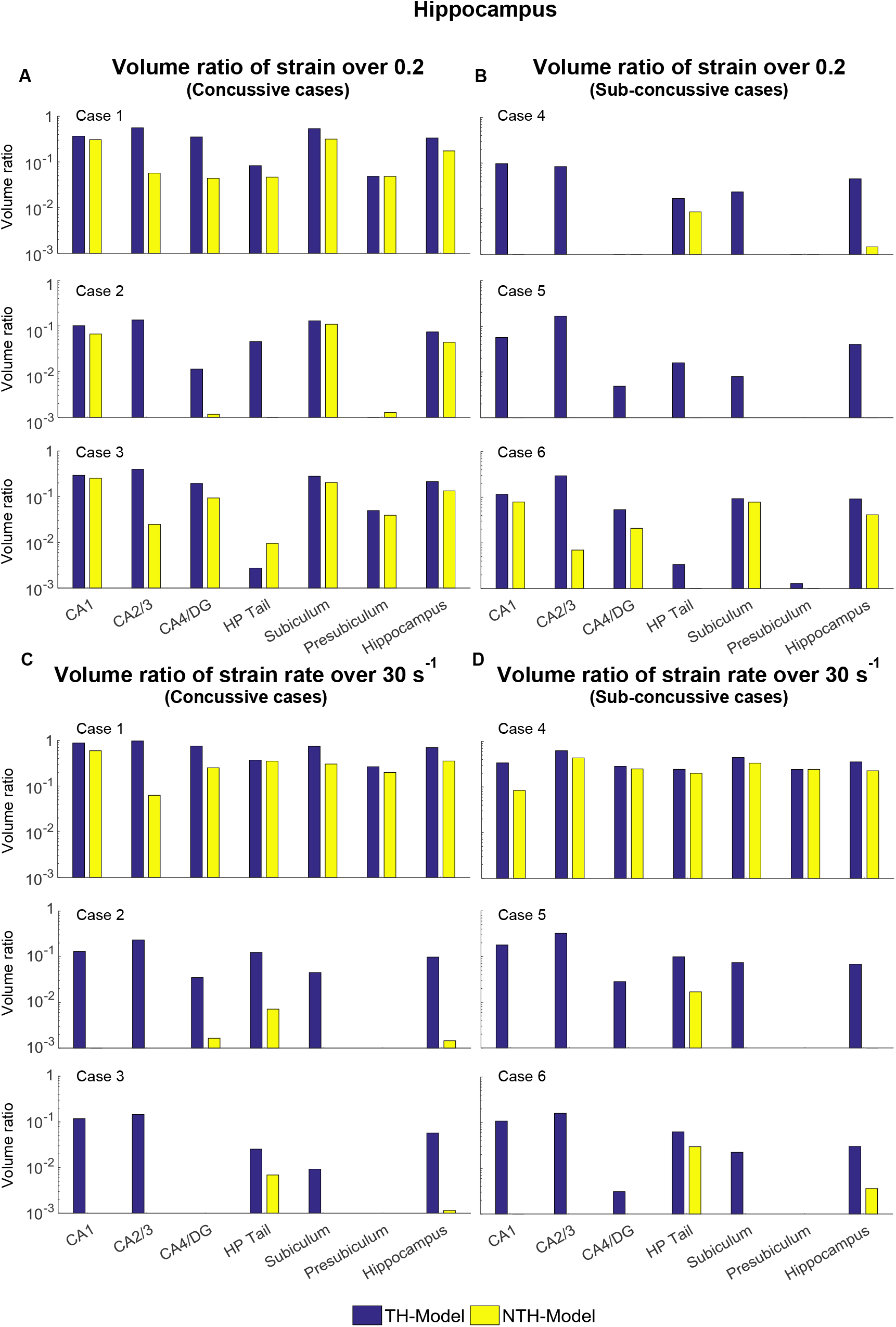
**Volume ratio of the maximum principle strain over 0.2 and strain rate over 30 s_-1_ in the hippocampal subfields and the whole hippocampus between the TH-Model and NTH-model of 3 concussive impacts (Cases 1-3) and 3 sub-concussive impacts (Cases 4-6).**

## Appendix E Comparison of strain and strain rate in the non-hippocampal regions

**Fig E1.**
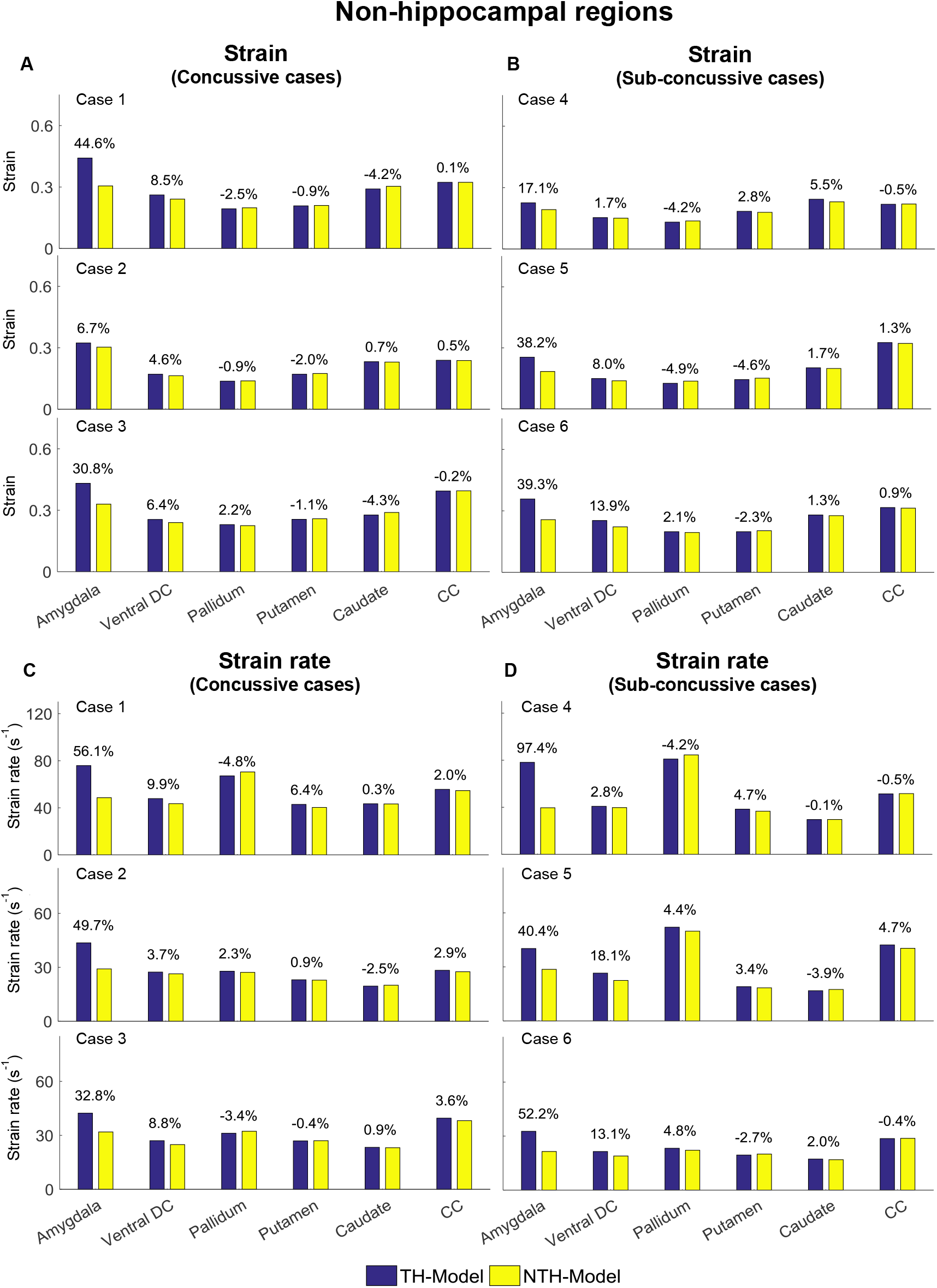
Comparison of strain and strain rate in the non-hippocampal regions between the TH-model and NTH-model of 3 concussive impacts (Cases 1-3) and 3 sub-concussive impacts (Cases 4-6). (A) Comparison of strain in the non-hippocampal periventricular regions of 3 concussive impacts. (B) Comparison of strain in the non-hippocampal periventricular regions of 3 sub-concussive impacts. (C) Comparison of strain rate in the non-hippocampal periventricular regions of 3 concussive impacts. (D) Comparison of strain rate in the non-hippocampal periventricular regions of 3 sub-concussive impacts. Percentages in strain difference and strain rate difference are calculated with the results of the NTH-model as the baseline. Ventral DC: ventral diencephalon; CC: Corpus callosum.

## Notes

### Competing Interest Statement

The authors have declared no competing interest.

